# Genome-Wide Mechanisms of Lifespan Extension by Dietary Restriction in Yeast

**DOI:** 10.1101/126797

**Authors:** Sergio E. Campos, Erika Garay, J. Abraham Avelar-Rivas, Alejandro Juárez-Reyes, Alexander DeLuna

## Abstract

Dietary restriction is arguably the most promising non-pharmacological intervention to extend human life and health span. Yet, only few genetic regulators mediating the cellular response to dietary restriction are known, and the question remains which other transcription factors and regulatory pathways are involved. To gain a comprehensive view of how lifespan extension under dietary restriction is elicited, we screened the chronological lifespan of most gene deletions of *Saccharomyces cerevisiae* under two dietary regimens, restricted and non-restricted. We identified 472 mutants with enhanced or diminished extension of lifespan by dietary restriction. Functional analysis of such dietary-restriction genes revealed novel processes underlying longevity specifically by dietary restriction. Importantly, this set of genes allowed us to generate a prioritized catalogue of transcription factors orchestrating the dietary-restriction response, which underscored the relevance of cell-cycle arrest control as a key mechanism of chronological longevity in yeast. We show that the transcription factor Ste12 is needed for full lifespan extension and cell-cycle arrest in response to nutrient limitation; linking the pheromone/invasive growth pathway with cell survivorship. Strikingly, *STE12* overexpression was sufficient to extend chronological lifespan under non-restricted conditions. Our global picture of the genetic players of longevity by dietary restriction highlights intricate regulatory cross-talks in aging cells.

## INTRODUCTION

Dietary restriction—a reduction in calorie intake without malnutrition, or substitution of the preferred carbon or nitrogen source—extends lifespan in virtually all species studied in the laboratory, from yeast to primates (Masoro 2005; Mair & Dillin 2008). Dietary restriction has been associated with protection against age-associated disease in mice, including neurodegenerative disorders (Zhu et al. 1999) and cancer (Yamaza et al. 2010), promoting not only a longer lifespan but also healthier aging (Fontana & Partridge 2015). Importantly, this intervention reduces the mortality rate in non-human primates (Colman et al. 2014), and delays the onset of aging-related physiological changes in humans (Holloszy & Fontana 2007), making dietary restriction the most promising intervention targeted to extend human lifespan. Yet, we are still missing a global picture of the genetic architecture of such lifespan response, which is needed to grant a deeper understanding of the genotype-phenotype relationship of aging and longevity (Schleit et al. 2013).

The budding yeast *Saccharomyces cerevisiae* has been a pivotal model organism in the discovery of the molecular and cellular basis of aging. Two aging models are widely used in this organism: The replicative lifespan of yeast, which refers to the number of times a single yeast cell can divide, and the chronological lifespan (CLS), which is a measure of the viability of a population during stationary phase throughout time. The latter provides a good model for inquiring the molecular changes and damage faced by post-mitotic cells (Longo et al. 2012).

In yeast, there is evidence that dietary restriction results in lifespan extension at least through the modulation of the conserved TOR and Ras/cAMP/PKA pathways, which regulate cellular growth and maintenance in response to nutrient availability (Kaeberlein et al. 2005). Depletion of TOR components, Tor1 and Sch9, results in CLS extension (Powers et al. 2006). Under low nutrient conditions, the serine/threonine kinase Rim15 phosphorylates transcription factors Msn2, Msn4, or Gis1, activating a maintenance response (Fabrizio et al. 2004). However, the *msn*2Δ*msn4*Δ*gis1*Δ triple mutant still shows lifespan extension by DR (Wei et al. 2008). Moreover, transcriptomic evidence and database analysis suggest that a larger number of up- and downstream genes are involved in lifespan extension (Wuttke et al. 2012; Choi et al. 2017); most of these candidates lack direct phenotypical confirmation. These observations suggest that there is yet to be identified an unknown number of regulators mediating lifespan extension by dietary restriction.

Research on aging in yeast has recently taken advantage of genome-wide approaches, enabling a comprehensive description of genes involved in lifespan regulation. For instance, a recent systematic study of replicative lifespan of most viable deletion strains revealed biological processes mediating longevity, such as translation, the SAGA complex, and the TCA cycle (McCormick et al. 2015). In the CLS model, several studies have aimed to estimate in parallel the stationary-phase survival of single-deletion mutants (Powers et al. 2006; Matecic et al. 2010; Fabrizio et al. 2010; Gresham et al. 2011; Garay et al. 2014), showing that autophagy, vacuolar protein sorting, regulation of translation, purine metabolism, chromatin remodeling, and the SWR1 complex are major determinants of stationary-phase survival. However, a direct comparison of the lifespan effects of such gene deletions under nutrient-rich and restricted conditions, that would allow to systematically address the mechanisms of longevity by dietary restriction is still missing.

The aim of this work was to generate a global picture of the underlying genetics of lifespan extension by dietary restriction, by systematically describing gene-diet interactions in yeast. Specifically, we compared the CLS of a collection of 3,718 knockout mutants aged under non-restricted and dietary-restricted media, using a high-resolution parallel phenotyping assay (Garay et al. 2014). These screens revealed 472 genes that influence the lifespan response to dietary restriction. Subsequent analyses uncovered the major biological processes and a comprehensive catalogue of transcription factors that control lifespan extension in a dietary-restriction regime. In particular, we revealed a link between Ste12 the pheromone/invasive growth transcription factor and lifespan regulation, suggesting that Ste12-mediated cell-cycle control is a mechanism of chronological longevity in response to nutrient limitation.

## RESULTS

### Systematic identification of dietary-restriction genes in yeast

We set out to describe, at the genome-wide level, the way in which dietary restriction influences the lifespan effect of gene mutations. In yeast, extended lifespan is achieved either by limiting the concentration of glucose in the growth medium (Koubova & Guarente 2003) or by using a non-preferred source of nitrogen (Powers et al. 2006; Jiang et al. 2000; Anderson et al. 2003). Here, dietary restriction was achieved by using GABA as the sole source of nitrogen, while non-restricted medium contained the preferred nitrogen-source glutamine (see Experimental Procedures). Dietary restriction increased half-life from 20.9 ± 0.4 days to 33.7 ± 1.3 days in the WT strain, which represents a 61% extension of the CLS (Figure 1A).

**Figure 1.**
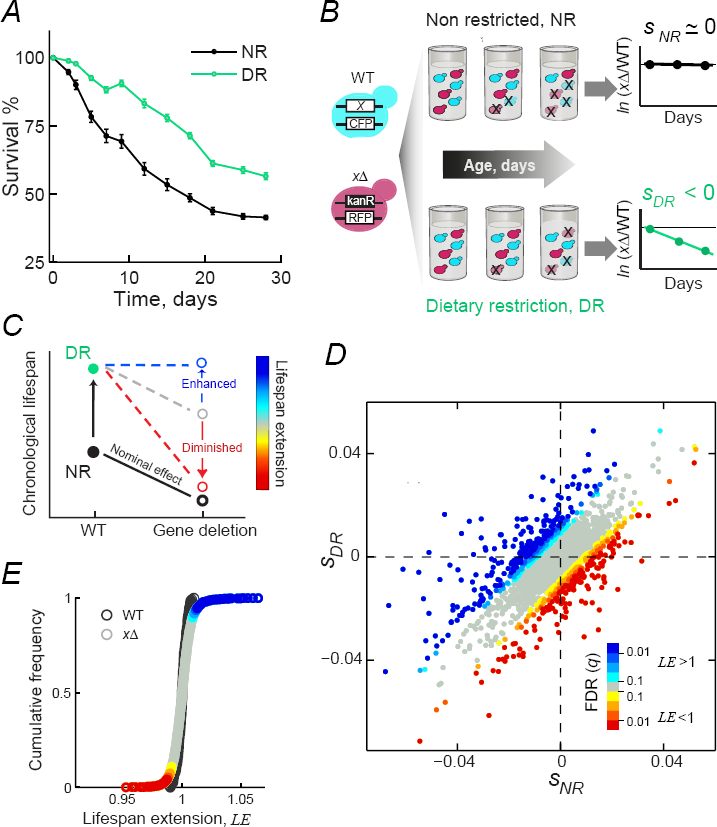
Genome-wide profiling of chronological-lifespan under two dietary regimes reveals a comprehensive compendium of dietary-restriction genes. (A) Survival curves of the WT strain under non-restricted (NR, black) and dietary restriction medium (DR, green). Error bars are the standard error of the mean (SEM, n=14). (B) Schematic representation of the genomic screening strategy; an RFP-labeled knockout strain (xΔ) is aged in co-culture with a CFP-labeled WT strain; a survival coefficient (s) is obtained for each aging condition. (C) Norm of reaction for possible outcomes of lifespan extension by dietary restriction; the detrimental effect of a short-lived knockout can be maintained (gray dashed line), aggravated (red dashed line), or alleviated (blue dashed line). (D) Scatter plot of the survival coefficients of 3,718 deletion strains under NR (horizontal axis) and DR (vertical axis); data points above or below the diagonal are colored according to the lifespan extension (LE) significance (q) compared to the WT replicates. (E) Cumulative distribution of *LE* values for mutant strains (circles) and WT replicates (crosses).

To identify the genetic determinants of lifespan extension by dietary restriction at the genome-wide level, we measured the CLS of 3,718 gene-knockout strains. We used a high-resolution profiling assay based on the measurement of a relative survival coefficient (s) of each knockout strain aged in co-culture with the WT strain (Garay et al. 2014) under dietary-restricted or non-restricted media (Figure 1B). The quantitative nature of our experimental data allowed us to determine whether the gene deletions had a neutral, deleterious, or beneficial effect on the CLS under each condition (Figure S1; Table S1). We scored 573 significantly short-lived and 254 long-lived single knockout strains in the non-restricted medium (FDR<5%), while dietary restriction resulted in 510 short-lived and 228 long-lived strains.

To validate our large-scale screen, we phenotyped a set of randomly-selected group of knockout strains that showed significant CLS effects. We used a previously-reported assay for quantitative analysis of yeast CLS (Murakami et al. 2008) to obtain viability curves for single-strain cultures as a function of time under the two nutrient conditions (Figure 2A–B). Twelve out of 16 (75%) strains re-tested under non-restricted medium recapitulated the CLS effects observed in the genome-wide screen, while 11 out of 17 (65%) strains tested were consistent with the results under dietary restriction (Figure S2; p<0.05, T-test). We note that this rate of false-positive hits is much lower than in other high-throughput assays that have used pooled deletion strains (Fabrizio et al. 2010; Matecic et al. 2010). Hence, our profiling approach provides accurate CLS scores for high-throughput profiling under different conditions, which allows scoring the gene-diet interactions influencing lifespan phenotypes.

**Figure 2.**
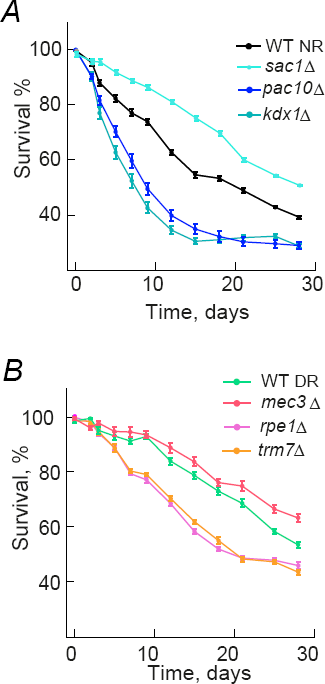
Validation of short and long-lived strains. Survival curves for selected gene knockouts that validated under (A) non-restricted conditions (NR) and (B) dietary restriction (DR). Survival curves and statistical analysis of the entire set of 16 (NR) and 17 (DR) strains tested is shown in Figure S2. Error bars are the S.E.M. (n=7).

To gain insight into the genes that mediate the lifespan-extending effects of dietary restriction, we searched for deletion strains that showed differential relative CLS effects when comparing the two diets (Figure 1C). Specifically, we looked for strains in which the mutant's lifespan relative to the WT was shorter in dietary restriction than in non-restricted medium (diminished lifespan extension). Likewise, we scored those cases in which the relative lifespan was relatively longer under dietary restriction (enhanced lifespan extension). In this way, by comparing the relative lifespans of all strains under non-restricted and dietary-restriction conditions, we were able to identify the set of gene knockouts leading to diminished or enhanced lifespan extension in yeast (Figure 1D).

We quantitatively defined the relative lifespan extension of each knockout as 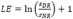 (see Experimental Procedures). The *LE* of each deletion strain was compared to the distribution of *LE* in 264 independent WT replicates to obtain a Z-score (Figure 1E). After filtering out strains that did not show a significant CLS effect in either condition, we obtained a list of 472 gene-knockouts with altered dietary-restriction response (FDR<5%; Table S2). This comprehensive set, which we termed DR-genes, includes 219 knockouts with diminished longevity (LE<1) and 253 gene-knockouts that displayed enhanced lifespan extension (LE>1). Together, these results show that many gene-environment interactions underlie longevity by dietary restriction in yeast.

### Functional classification of dietary-restriction genes

To describe which downstream cellular functions influence lifespan extension by dietary restriction, we sought to classify the 472 DR-genes according to their annotated functional features. We used a kappa statistic approach (Huang et al. 2009) to cluster genes by shared GO terms and mutant phenotypes, as reported in the Saccharomyces Genome Database (see Experimental Procedures). The analysis was performed separately for genes with diminished (LE<1) or enhanced (LE>1) lifespan extension (Figure 3; Table S3). Some clusters recapitulated cellular functions previously related to lifespan regulation, such as autophagy (Meléndez et al. 2003), mitochondrial function (Ocampo et al. 2012; Aerts et al. 2009), and cytosolic translation (Pan et al. 2007; Hansen et al. 2007). Importantly, this classification also identified novel processes such as maintenance of cell-cycle arrest mediated by pheromone and establishment of nucleus localization in the cell. These observations show that our screen was able to independently identify processes that were previously known to influence the dietary response and at the same time allowed the identification of novel genes and processes related to longevity by dietary restriction.

**Figure 3.**
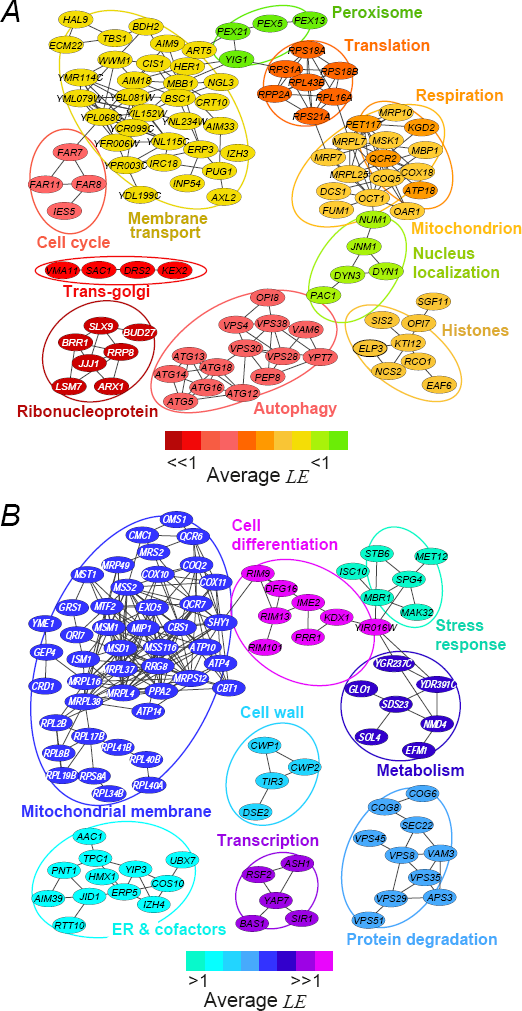
Functional clustering of dietary-restriction genes. Network representation of functional clusters of genes with (A) LE<1 and (B) LE>1. Edges denote agreement between pair of genes *(kappa>0.35).* Node color code indicates decreasing or increasing ranks of mean *LE* of the cluster.

Deletion of genes necessary for the maintenance of cell-cycle arrest resulted in a diminished lifespan extension. Specifically, deletion of pheromone-responsive genes *FAR7* and *FAR8* had short-lived phenotype under dietary restriction (Figure 3A). In yeast, these Far proteins prevent cell cycle recovery after pheromone exposure, possibly by inhibiting *CLN1-3* (Kemp & Sprague 2003). Moreover, mutations in genes required for processing and correct localization of ribonucleoprotein complexes resulted in a strongly diminished lifespan extension. While it is well established that ribosomal function is downregulated in response to TOR inhibition or nutrient depletion (Hansen et al. 2007), our screen pointed to specific proteins involved in pre-rRNA processing (Slx9), nucleolar rRNA methyltransferase (Rrp8), nuclear export of pre-ribosomal subunits (Arx1), and translational initiation (Bud27). Likewise, deletion of *DYN1-3, JNM1, NUM1,* and *PAC1,* all involved in nuclear movement along microtubules, resulted in diminished lifespan extension, suggesting that microtubule dynamics underlies dietary restriction. In this regard, it is known that certain cellular processes needed for extended longevity, such as autophagy, require intact function of microtubules (Köchl et al. 2006). However, the relationship between nuclear localization and lifespan extension remains unexplored.

We also found clusters with enhanced lifespan extension, such as mitochondrial function (Figure 3B). While dietary restriction shifts the metabolism towards respiration (Lin et al. 2002), impaired respiration promotes longevity in yeast and nematodes through enhanced retrograde response and activation of anaplerotic pathways (Cristina et al. 2009). Hence, a higher demand for respiration during dietary restriction could lead to the activation of compensatory pathways; similar feedback mechanisms might account for the alleviation of deleterious effects observed in other deletion strains. For example, the short-lived phenotypes of deletion of cell-wall genes *CWP1, DSE2,* and *TIR3* were largely alleviated under dietary restriction. Taken together, these findings suggest that lifespan extension in response to dietary restriction in yeast is a complex phenotype resulting from the interplay of many downstream cellular processes.

### A defined set of transcription factors regulate lifespan extension by dietary restriction

Complex phenotypic responses are frequently coordinated by transcriptional regulation of functionally-related genes. To investigate the transcriptional regulation of longevity by dietary restriction, we analyzed our set of DR-genes using an algorithm to search for the transcriptional regulators of these genes. We used TFrank (Gonçalves et al. 2011), a graph-based approach that takes advantage of available interactions of transcription factors and their targets, to obtain a list of prioritized regulatory players within the yeast regulatory network (Table 1). This approach allowed us to assess the possible lifespan role of transcription factors that were not included in our genome-wide screen due to gene essentiality or sterility. Transcription factors within the top 5% rank of this analysis included Msn2 and Msn4; these transcription factors are well known regulators of lifespan (Fabrizio et al. 2004). However, most top-ranked transcription factors had not been previously associated to lifespan extension by dietary restriction in yeast. Noteworthy, the top hits (Ace2, Ash1, Tec1, Spf1, and Ste12) are known to regulate different aspects of cell-cycle progression.

**Table 1.** A catalogue of transcription factors regulating dietary-restriction genes in yeast (top 5% rank)

| Rank | TF | Weight<sup>a</sup> | % Regulated<sup>b</sup> | Description |
| --- | --- | --- | --- | --- |
| 1 | Ace2 | 3.15 | 75.5 | Involved in G1/S transition of the mitotic cell cycle; activates cytokinetic cell separation |
| 2 | Ash1 | 2.94 | 50.4 | Negatively regulates G1/S transition of mitotic cell cycle; activates pseudohyphal growth |
| 3 | Tec1 | 2.81 | 63.0 | Transcription factor targeting pseudohyphal-growth genes and Ty1 expression |
| 4 | Sfp1 | 2.55 | 66.3 | Regulates ribosomal-protein genes, response to nutrients and stress, and G2/M transitions of the mitotic cell cycle |
| 5 | Ste12 | 2.45 | 58.2 | Activates genes involved in mating or pseudohyphal-growth pathways |
| 6 | Bas1 | 2.36 | 44.9 | Involved in regulating the expression of genes of purine and histidine biosynthesis |
| 7 | Snf6 | 2.19 | 37.4 | Subunit of the SWI/SNF chromatin remodeling complex |
| 8 | Msn2 | 2.10 | 54.3 | Regulation of transcription in response to a wide variety of stresses |
| 9 | Yrm1 | 1.79 | 44.5 | Transcription factor involved in multidrug resistance |
| 10 | Gcn4 | 1.76 | 46.7 | Activator of amino acid biosynthetic genes; responds to amino acid starvation |
| 11 | Ixr1 | 1.73 | 26.4 | Transcriptional repressor that regulates hypoxic genes during normoxia |
| 12 | Abf1 | 1.723 | 45.3 | DNA binding protein with possible chromatin-reorganizing activity |
| 13 | Msn4 | 1.65 | 44.0 | Regulation of transcription in response to a wide variety of stresses |
| 14 | Rap1 | 1.53 | 41.4 | Essential DNA-binding transcription regulator; role in chromatin silencing and telomere length |

**a.** TFRank weight (Gonçalves et al. 2011).
**b.** Percentage of dietary-restriction genes regulated by the transcription factor.

To directly confirm the role in lifespan regulation of TFRank hits, we generated *de novo* deletion mutants for seven transcription factors and characterized their CLS under non-restricted and restricted conditions (Figure 4). In agreement with the *in silico* inference, deletions of six transcription factors resulted in short-lifespan under dietary restriction, while the reminder strain (*ace2*Δ) was long-lived. Most of the observed lifespan effects were moderate, which is in agreement with the fact that transcription factors in yeast typically act upon overlapping targets, providing functional compensation to one another (Zheng et al. 2010). These results show that high-TFRank hits are regulators of lifespan extension by dietary restriction in yeast.

**Figure 4.**
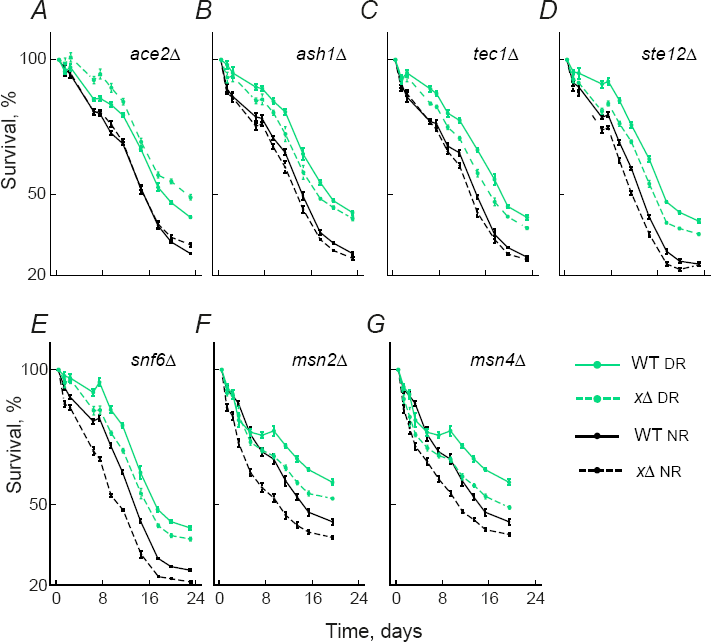
Deletion of transcription factors genes that regulate dietary-restriction genes affects CLS. Survival curves of gene deletions and WT strains aged under non-restricted medium (NR, solid black line) or dietary restriction (DR, solid green line). Deletion strains (black or green discontinuous lines, for NR or DR, respectively) are for genes coding for transcription factors Ace2 (A), Ash1 (B), Tec1 (C), Ste12 (D), Snf6 (E), Msn2 (F), and Msn4 (G). Error bars are the S.E.M. (n=7).

### Ste12 is a positive regulator of longevity by dietary restriction and cell-cycle arrest in response to nutrients

Among the top hits of our transcription-factor analysis was Ste12, which acts on downstream genes involved in mating or pseudohyphal growth (Dolan et al. 1989; Roberts & Fink 1994). To further explore the regulatory role of Ste12 in lifespan extension, we characterized the CLS of the *ste12*Δ strain under different concentrations of glucose. Lifespan extension was partially abrogated in the *ste12*Δ strain under glucose restriction and, to a lesser extent, under high concentration of glucose (Figure 5A). We also characterized the CLS of this strain using standard aerated culture conditions (Hu et al. 2013), which unambiguously confirmed that Ste12 is a positive regulator of lifespan extension in yeast (Figure S3).

**Figure 5.**
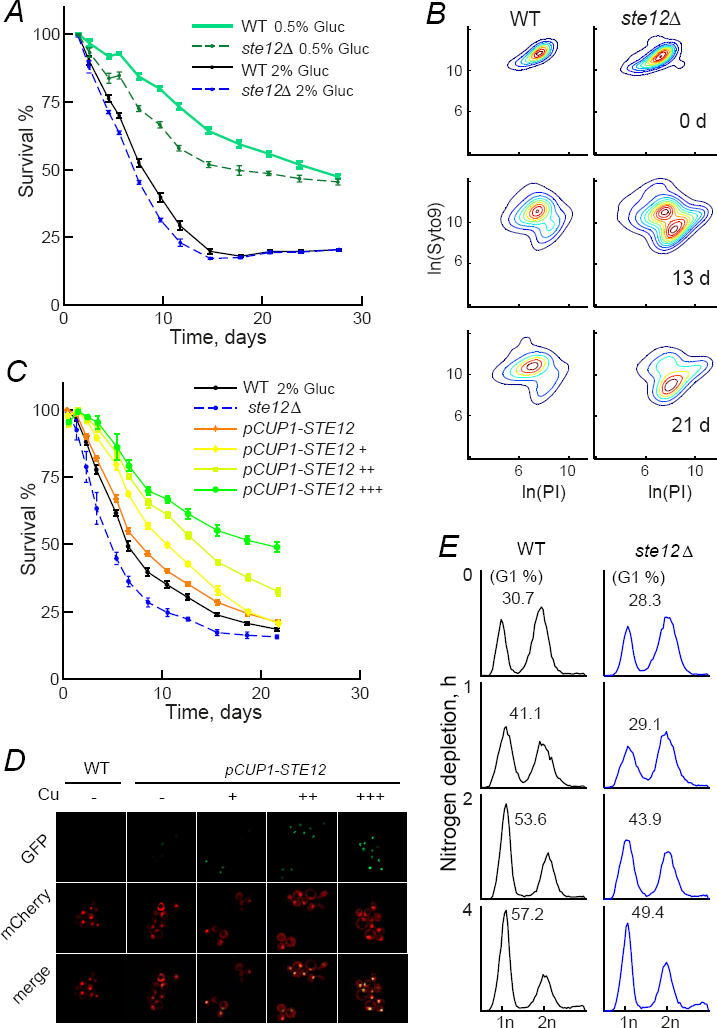
Ste12 is a positive regulator of lifespan extension by dietary restriction. (A) Survival curves of WT and *ste12Δ* strains aged in 0.5% or 2% glucose. Error bars represent S.E.M. (n=7). (B) Contour plots showing WT and *ste12Δ* populations of dead and alive cells under SC 0.5% glucose; fluorescence of Syto9 (alive cells) and propidium iodide (PI, dead cells) is shown in the vertical and horizontal axes, respectively, across time in stationary phase. (C) Survival curves of the WT, *ste12Δ*, and *pCUP1::GFP-STE12* overexpression strains aged in SC 2% glucose in non-induced conditions (orange) or induced with 2μM (+), 5μM (++), or 15 μM (+++) copper sulfate. Error bars are the S.E.M. (n=7). (D) Micrographs of WT and *pCUP1::GFP-STE12* strains with GFP, mCherry and merge filters, showing induction with varying copper sulfate concentrations. (E) DNA content labeled with SYTOX Green and measured by flow cytometry; fluorescence of DNA *vs.* normalized cell count is shown for WT and *ste12Δ* strains challenged with a nitrogen depletion shock for 0, 1, 2, or 4 hours. Numbers indicate percentage of cells in the G1 cell-cycle phase.

Most methods for measuring CLS in yeast—including ours—rely in the ability of stationary-phase cells to re-enter the cell cycle upon transfer to fresh medium. Given that Ste12 is involved in cell-cycle arrest, we decided to rule out any possible methodological artifact by directly measuring the fraction of dead relative to alive strains in stationary phase. This experiment showed that, under limited glucose concentration, the alive *ste12*Δ population decayed faster than the WT under calorie restriction (Figure 5B; Figure S3). These results indicated that the CLS-effects of the *STE12* deletion are maintained regardless of the methodology and nutrient restriction condition used to infer population survivorship in stationary phase.

A bona fide positive regulator of lifespan is expected to increase cell survivorship when over-expressed. We thus investigated whether high *STE12* expression causes lifespan extension under non-restricted conditions. To this end, we generated a copper-inducible strain with a GFP fusion to track Ste12 protein levels. The WT and *pCUP1-STE12* strains were aged under varying concentrations of copper sulfate in 2% glucose SC non-restricted medium. We found that the CLS of the STE12-overexpression strain increased readily as a function of copper concentration (Figure 5C), while in the WT strain increasing copper concentrations had no significant effect on CLS nor growth (data not shown). Importantly, GFP signal in the nucleus also increased as a function of copper concentration, confirming that lifespan extension was linked to increased levels of Ste12 in the nucleus (Figure 4D). Together, these results indicate that Ste12 is a novel positive transcriptional regulator of lifespan in yeast.

Transcriptional targets of Ste12 are known to mediate G1 arrest of the cell-cycle in response to pheromone signal (Dolan & Fields 1990), but the role of such Ste12-mediated cell-cycle arrest in aging is unknown. We asked whether Ste12-mediated cell-cycle control in response to nutrient limitation is a cellular mechanism of lifespan extension by dietary restriction in yeast. To this end, we used flow cytometry to monitor the cell-cycle dynamics of yeast populations following nitrogen starvation (Figure 4E). Our results show that *ste12*Δ cells failed to arrest the cell cycle at the same rate as the WT strain. This observation suggests that the transcription factor Ste12 integrates nutrient signaling and regulates downstream genes needed for cell-cycle arrest, which may underlie its beneficial effect on cell survivorship.

### Ste12 mediates lifespan extension in association to Tec1

To gain further insight on the cellular pathways of lifespan extension by dietary restriction, we asked whether the lifespan effect of *STE12* was linked to Tec1, a transcription factor of the invasive-growth pathway. Ste12 and Tec1 form a heterodimer that activates filamentous-growth genes (Madhani et al. 1999). Moreover, Tec1 has also been associated to lifespan regulation in response to rapamycin (Brückner et al. 2011). We generated a *tec1*Δ*ste12*Δ double-deletion mutant and found an epistatic interaction between these genes under glucose restriction; the CLS phenotypes of the *ste12*Δ*tec1*Δ double deletion was not the additive combination of the single-deletion phenotypes (Figure 6A). This genetic interaction suggests that Tec1 acts in concert with Ste12 to promote longevity by dietary restriction.

**Figure 6.**
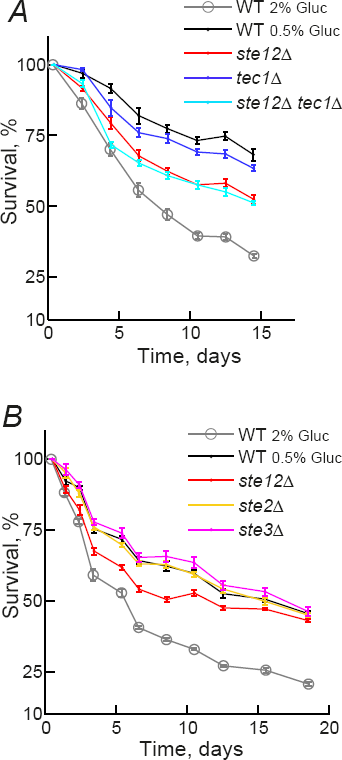
Ste12 promotes longevity in association to Tec1, but not to the pheromone receptors. (A) Survival curves of *ste12Δ* and *tec1A* single and double knockout strains under dietary restriction. (B) Survival curves of *ste2Δ* and *ste3Δ* single knockouts under dietary restriction.

Finally, we asked whether the effect of Ste12 is associated to elements of the pheromone-responsive MAPK pathway. Deletion of the pheromone-receptor genes *STE2* and *STE3* did not show a lifespan-extension defect in response to glucose restriction (Figure 6B). Altogether, these results indicated that Ste12 regulates lifespan extension in response to nutrients through functional association to Tec1 and the invasive-growth pathway, but not under the control of upstream pheromone-pathway components.

## DISCUSSION

"The intrinsic nature of the ageing process is essentially one of systems degradation" (Kirkwood 2008). With a growing number of genetic aging factors in hand, the next great challenge is to understand how the different mechanisms underlying aging and longevity are integrated to one another and to the environment. In this study, we have adapted the chronological-aging paradigm in yeast to provide a quantitative and systematic description of how different genetic and dietary factors influence lifespan. Specifically, we have screened a collection of 3,718 gene-deletion strains for short and long-lived phenotypes under two dietary regimes. Our analysis revealed 472 genes (DR-genes) in which dietary restriction modifies the relative lifespan effect of the gene knockout. To the best of our knowledge, this study yields the most comprehensive phenotypic compendium of genetic players involved in longevity by dietary restriction.

The downstream genetic players involved in longevity by dietary restriction included autophagy, cytosolic translation, peroxisome biogenesis, respiration and mitochondrial function, along with novel factors such as the cell-cycle arrest machinery, processing of ribonucleoprotein complexes, cell-wall organization, and microtubule-based nuclear movement. Furthermore, our analysis of DR-genes as regulatory targets allowed the establishment of a set of ranked transcription factors that orchestrate the cellular response to dietary restriction (Table 1). Among the top hits were Msn2 and Msn4, two positive regulators of stress-response and lifespan extension which operate downstream of the Tor/Sch9/Ras-PKA pathways that converge on Rim15, the main protein kinase involved in cell survivorship in response to nutrients (Wei et al. 2008). Strikingly, high-ranked transcription factors were regulators of mitotic cell-cycle transitions by either repression of Cln3 specifically in yeast daughter cells (Ace2 and Ash1) (Laabs et al. 2003; Di Talia et al. 2009), activation of ribosomal-protein genes (Sfp1) (Marion et al. 2004), or cell differentiation in response to nutrients or pheromone (Tec1 and Ste12) (Madhani et al. 1999). This observation underscores the role of cell-cycle control as a key mechanism of chronological lifespan in yeast, providing a defined set of transcription factors that could act downstream of Rim15, which plays a crucial role in cell-cycle arrest upon nutrient limitation (Weinberger et al. 2007).

We have provided compelling evidence that *STE12* is a positive regulator of longevity by dietary restriction. Not only did the *ste12*Δ strain show diminished extension by dietary restriction measured both by outgrowth and dead/alive assays, but also *STE12* overexpression was sufficient to extend lifespan under a non-restricted diet. Importantly, the *STE12* phenotypes in our screening settings were confirmed under standard chronological-aging conditions. Ste12 is a transcription factor downstream of two cell-differentiation programs regulated by MAPK pathways, namely mating and invasive growth (Dolan et al. 1989; Madhani et al. 1999). We also found that deletion of *STE12* results in a failure to arrest the cell cycle upon nutrient starvation. In the presence of pheromone, the cell cycle is arrested by the action of Ste12, Far1, and the FAR complex (Far3 and Far7-11) (Kemp & Sprague 2003); Far1 and Far3 are direct targets of Ste12 (Lefrançois et al. 2009). Since efficient G1 arrest protects against replication stress during stationary phase, effectively extending CLS (Weinberger et al. 2007), we propose that cell-cycle control underlies the lifespan effect of Ste12 through control of the FAR proteins. Concomitantly, deletion of *FAR7, FAR8,* and *FAR11* in our screen resulted in diminished lifespan extension. It is thus likely that regulatory elements of the mating/pheromone pathway are recruited to arrest the cell cycle during dietary restriction.

We have shown that deletions of *STE12* and *TEC1* interact in an epistatic manner, which suggests that these transcription factors act in concert to promote yeast lifespan. Ste12 is associated to Tec1 during pseudohyphal growth (Chou et al. 2006). In addition to controlling cellular development in response to stimuli, Tec1 is needed for full lifespan extension in response to the Tor1-inhibiting drug rapamycin (Brückner et al. 2011), suggesting that Tec1 acts downstream of the TOR pathway. Also, transcription analyses have shown that Ste12 is a regulatory hub during stationary phase under rapamycin treatment (Wanichthanarak et al. 2015), further strengthening the idea that Ste12 and Tec1 link TOR and MAPK-signaling pathways. However, Tec1 promotes cell-cycle progression by activation of Cln1 (Madhani et al. 1999), in conflict with the fact that the cell-cycle is arrested in response to nutrient limitation. It is thus likely that the players upstream Ste12 and Tec1 are involved in an intricate signaling response leading to lifespan extension.

It remains to be addressed whether the role on aging of the set of transcription factors identified in this study is conserved. For instance, *STE12* has no clear homolog in animals, however, transcriptional networks can be rewired through evolution, leading to changes in the regulation exerted by specific regulators, while the downstream targets remain associated (Sorrells et al. 2015). In addition, the yeast three-kinase module regulating MAPK pheromone and invasive-growth pathways are conserved in other organisms (Widmann et al. 1999). In particular, *KSS1* and *FUS3* are key members of the MAPK pathway that regulates cell differentiation programs in yeast (Bardwell 2004), while their mammalian counterpart *MAPK1* has been reported to regulate cell-fate determination (Chaman et al. 2015). Also, the MAPK1/ERK pathway is central to the development of several age-associated diseases in mammals (Carlson et al. 2008). Thus, the study of targets downstream the MAPK pathway in yeast might bring important insights into the regulation of aging in other eukaryotes, including humans.

Our genome-wide screens provide a much-needed comprehensive picture of the mechanisms of lifespan extension by dietary restriction, underscoring links between nutrient sensing, the cell-cycle arrest machinery, and longevity in yeast. Our approach can be readily applied to other genetic, environmental, or pharmacological perturbations, providing a systematic framework to describe aging networks in a simple tractable system. Other cross-talks among downstream cellular processes and their transcriptional regulators may remain to be uncovered, which will shed further light to the genetic wiring of aging cells.

## EXPERIMENTAL PROCEDURES

### Strains and media

Fluorescent single-gene deletion strains are prototrophic haploids (*MATa xxx*Δ*::kanMX4 PDC1-mcherry-CaURA3MX4 can1*Δ*:STE2pr-SpHIS5 lyp1*Δ *his3*Δ*1 ura3*Δ*0 LEU2*) derived from crossing the *MATα* YEG01-RFP SGA-starter with strains from the MATa BY4741 deletion collection, as previously described (Garay et al. 2014). All *de novo* single-gene deletions were generated in the YEG01-RFP parental strain *(MATα PDC1-mcherry-CaURA3MX4 can1*Δ*:STE2pr-SpHIS5 lyp1*Δ *his3*Δ*1 ura3*Δ*0 LEU2)* by direct gene replacement with the *natMX4* module conferring resistance to clonNAT; double-knockouts were generated from such *de novo* single-gene deletions using the hygromycin-resistance *hphMX4* cassette. The Ste12 overexpression strain was generated by inserting the *CUP1* promoter and GFP-fusion construct from plasmid pYM-N4 in the 5’ region of *STE12* ORF, as described (Janke et al. 2004).

Non-restricted (NR) aging medium contained 0.17% yeast nitrogen base (YNB) without amino acids and ammonium sulfate, 2% glucose, 0.07% amino acid supplement mix (Cold Spring Harbor Laboratory Manual 2005), and 25 mM glutamine as nitrogen source. Dietary-restricted (DR) aging medium was prepared substituting glutamine with 25 mM of γ-amino butyric acid (GABA). The choice of a non-preferred nitrogen source for DR instead of limited glucose concentrations overcomes the dramatic metabolic changes due to glucose repression in yeast (Kresnowati et al. 2006), while facilitating the parallel characterization of stationary-phase cultures in low volumes, given that yields are similar under NR and DR conditions. SC medium used for DR based on glucose concentration was 0.17% yeast nitrogen base (YNB) without amino acids, 0.5% or 2% glucose, and 0.2% amino acid supplement mix.

All outgrowth cultures were performed in low-fluorescence medium (YNB-lf). Nitrogen-starvation medium for cell-cycle progression experiments was 2% glucose and 0.17% YNB without amino acids and ammonium sulfate.

### Automated competition-based CLS screens and data analysis

Fresh cultures of the RFP-tagged gene-deletion collection were replicated in 96-well plates (Corning 3585) with 150 μl of NR or DR aging medium. Saturated cultures were mixed with a CFP-labeled WT reference strain in a 2RFP:1CFP ratio, replicated by pinning into 96-deepwell plates (Nunc 260251) containing 700 μl of NR or DR aging medium, and grown at 30°C and 70% relative humidity, without shaking, in an automated cell-assay system (Tecan Freedom EVO200) integrated to a multi-label plate reader (Tecan M1000). We have previously shown that the CLS effects of mutants aged under low aeration in low-volume cultures change minimally compared to highly-aerated cultures (Garay et al. 2014). Data acquisition and initial processing have been described. In brief, five days after inoculation, 5 μl outgrowth cultures were inoculated every other day into 150 μl of fresh low-fluorescence medium; absorbance at 600nm (OD_600_) and fluorescence *(RFP* and *CFP)* measurements were taken every 150 min throughout 14 hrs. An apparent survival coefficient, *s*, and its standard error, *σ_s_,* were obtained from the slope of the linear regression (Robustfit, Matlab) of the log of the ratio of *RFP* to *CFP* signal at a fixed interpolation time point in the outgrowth culture (10 hrs), and the number of days in stationary phase, as previously described (Garay et al. 2014).

### Scoring CLS phenotypes and lifespan extension coefficients

Short- and long-lived knockouts under NR or DR were determined by assigning a *Z*-score to each mutant's s coefficient; the distribution's mean and standard deviation were from the measurement of 264 WT_RFP_/WT_CFP_ independent co-cultures under either condition. Two-tailed *p*-values were obtained from each *Z*-score to compute a false-discovery rate (FDR); we assigned significant phenotypes using a q<0.05 cutoff.

The effect of dietary restriction on the gene-deletion strains relative to the WT was evaluated by calculating their relative lifespan extension defined as 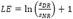, where *s_NR_* and *S_DR_* are the s coefficients of a given deletion strain obtained from the screen under NR and DR, respectively. A Z-score was assigned to the *LE* of each mutant compared to the distribution of *LE* values of 264 independent WT reference experiments, and this was used to obtain an FDR, significant *LE*<1 and *LE>*1 values were assigned using a q<0.05 cutoff.

### Survival curves of individual stationary-phase populations

Selected strains were grown individually in NR or DR aging medium for 48 hours at 30°C 200 rpm in aerated tubes, then transferred to 96-well plates. These plates were replicated onto 96 deep-well plates containing 700 μl of NR or DR medium and left for the entire experiment at 30°C and 70% relative humidity without shaking. From here on, all experimental steps were performed in an automated robotic station (Tecan Freedom EVO200). After 4 days, 10 μl aliquots were taken with an automated 96-channel pipetting arm to inoculate 96-well plates containing 150 μl of low fluorescence medium. OD_600_ was obtained in a plate reader (Tecan M1000) every 1.5 hours until saturation was reached; this first outgrowth curve was regarded as the first time point (T_0_, age = 0 days). Sampling was repeated every 2-3 days for 24-28 days. From these outgrowth curves, we extracted the doubling time and the time shift to reach mid-exponential phase (OD_600_=0.3) that occurred between the first day of measurements (*T*_0_) and each day in stationary phase (*T_n_*). Relative cell viability was calculated from these data, as reported by Murakami *et al* (Murakami et al. 2008).

Viability data points relative to T_0_ were used to plot a survival curve, which was fitted to an exponential decay model *(N(T) = N_0_e^−rT^*) where *N_0_* is the percentage of viability at *T*_0_, *T* is time in days, and *r* is the rate of death. For validation of CLS effects, mutants were taken from the RFP-tagged deletion collection (or generated *de novo,* when indicated) and viability was assayed to calculate death rates in at least 7 experimental replicates, which were compared to replicates of a WT strain; significant CLS effects were considered using a *p*<0.05 cutoff (*T*-test).

### CLS assay in standard aeration conditions

Pre-inoculums from three different colonies of each strain were set in 5 mL SC medium for 24 hours, these were diluted 1:200 v/v in 10 mL SC with 2% glucose or 0.5% glucose aging medium in 50 mL tubes for 48 hours with shaking (200 rpm) at 30 °C. Viability at each measuring point was obtained by monitoring the change in outgrowth-curve parameters with time in stationary phase, as described above.

### Visualization of functional clusters

Gene Ontology (GO) associations and phenotype terms were downloaded from the *Saccharomyces* Genome Database (SGD, last updated December 2016) to build two *m* by *n* matrixes, where *m* is the number of DR genes (219 and 253 for LE<1 and LE<1, respectively) and *n* is the number of GO and phenotypic terms (1,748). Each term was used to evaluate the overall agreement between gene-pairs to calculate Cohen’s *kappa* (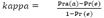 for each gene pair, where Pra(a) is the relative observed agreement or the number of terms that a gene-pair shares divided by the total number of terms in the matrix, and *Pr(e)* is the probability of agreement by chance, calculated as the sum of probabilities for each member of the gene-pair to be associated or not to each term.

Kappa values were used to build a matrix that represented the agreement between each gene-pair. Gene-pairs that showed *kappa>0.35* were regarded as likely similar and thus used as cluster seeds to form larger groups of genes; groups sharing more than 50% of their genes were merged in subsequent iterative steps. Clusters with at least four elements were manually named by inspection in the SGD and GO enrichment. Network representation was created using Cytoscape; edges between nodes represent kappa agreement above the established threshold (*kappa*>0.35).

### Alive/dead staining assay

We used the same scheme of 96 deep-well plates (one plate per replicate) to age cells in SC with 0.5% or 2% glucose. Each day, a single well of each strain was collected. Cells were centrifuged, washed, and dyed with LIVE/DEAD® FungaLigth™ Yeast Viability Kit, following manufacturer's instructions. Propidium iodide (IP) and Syto®9 fluorescence were measured by cell cytometry (LSRFortessa™, Becton Dickinson) at early stationary phase (4 days after inoculation) and at different time-points until 21 days in stationary phase. IP was excited with a 591-nm laser, fluorescence was collected through a 586/15 band-pass filter; Syto9 was excited with a 488-nm laser, and fluorescence was collected through 525/50 band-pass and 505LP emission filters. Cell-viability percentage was obtained from the total amount of cells measured and subtracting the number of dead-cell events.

### Cell-cycle assay

WT and mutant strains were grown in flasks containing 50 ml of NR aging medium at 30°C and shaken at 200 rpm until mid-logarithmic phase (OD_600_≅0.5). Cells were centrifuged, washed twice with sterilized water, and transferred to nitrogen starvation medium. Samples were taken at the moment of transfer (time 0) and 1, 2 and 4 hours after this time point. Fixation and dying with SYTOX™ Green were performed as described elsewhere. Cells were analyzed by flow cytometry (FACSCalibur™, Becton Dickinson); SYTOX™ Green was excited with a 488-nm laser, and fluorescence was collected through a 525/50 band-pass filter.

## Acknowledgements

We are grateful to Cei Abreu-Goodger, Tobias Bollenbach, and Eugenio Mancera for critical reading of the manuscript, and to Selene Herrera for technical assistance.

### Funding

This work was funded by the Consejo Nacional de Ciencia y Tecnología de México (CONACYT grant CB2015/164889), the University of California Institute for Mexico and the United States (UC MEXUS-COANCYT grant CN15-48), and the Consejo de Investigación sobre Salud y Cerveza de México. SEC had a doctoral fellowship from CONACYT. The funders had no role in study design, data collection and analysis, decision to publish, or preparation of the manuscript.

### Author contributions

SEC, EG, and AD conceived and designed the experiments. SEC, EG, JAA-R, and AJ-R performed the experiments. SEC and AD analyzed the data. SEC and AD wrote the paper. All authors read and approved the final manuscript.

## SUPPORTING INFORMATION LISTING

**Fig. S1.** Chronological-lifespan effects from genome-wide screens under two dietary regimes.

**Fig. S2.** Validation of hits from the genome-wide screens.

**Fig. S3.** STE12 deletion displays diminished lifespan extension measured by standard aeration conditions and alive/dead staining.

**Table S1.** Genome-wide CLS screens under two dietary regimes, complete data set (XLS).

**Table S2.** Data set of 472 DR-genes identified in this study (XLS).

**Table S3.** Functional clusters of DR-genes with diminished or enhanced relative lifespan extension (XLS).

---

**Figure S1.**
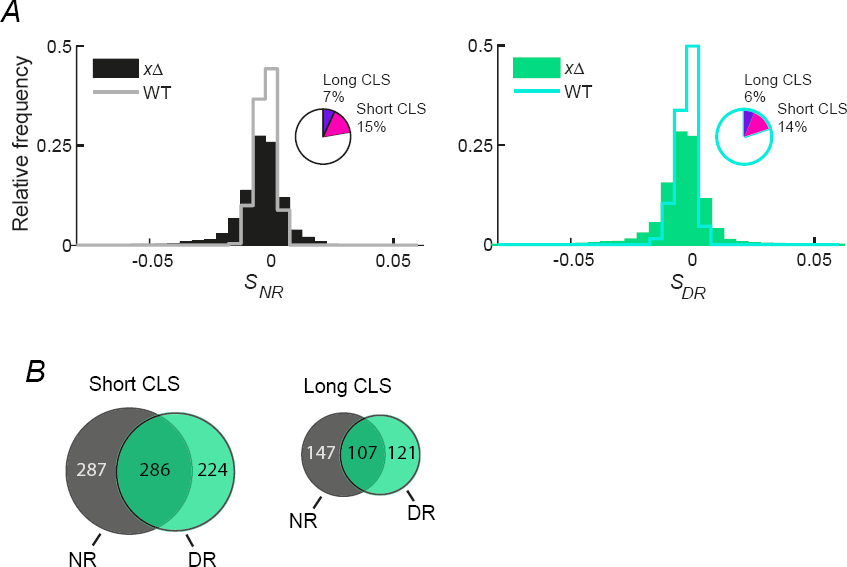
CLS effects from two genome-wide screens. (A) Histograms of the normalized counts for 3,719 single-knockout strains under non-restricted medium (NR, black) and dietary restriction (DR, green); WT controls (264) aged under NR (gray line) and DR (light green line) are shown. A distribution of the s coefficients of the WT-strain replicates was used to calculate a Z-score for each knockout strain and to define short-lived and long-lived strains (see Materials and Methods). Insets show the fraction of short-lived (magenta) and long-lived (purple) deletion strains scored in each screen. (B) Venn diagrams show the common number of short- and long-lived knockout strains scored under non-restricted medium (NR, dark gray) and dietary restriction (DR, green).

**Figure S2.**
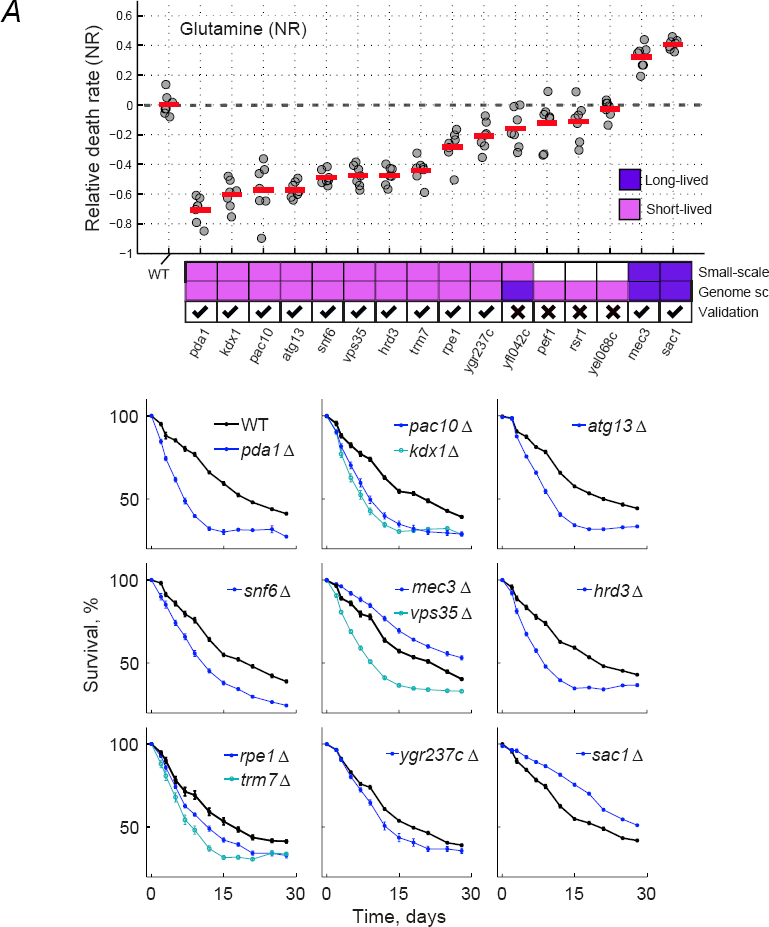
Validation of hits from the genome-wide screens. Top plot shows the relative death rates of WT and mutant strains, -1·/*n*(*r_x_/r_WT_*), aged under (**A**) non-restricted (Glutamine) and (**B**) dietary restriction (GABA) media. Death rates are from the fit of each survival curve to an exponential decay model. A CLS phenotype from the genome-wide screen was considered to be validated when the death rate of the mutant was consistent with the screen (short- or long-lived) and significantly different from that of the WT (*p*<0.05, two-tailed *T*-test). Bottom panels show the survival curves of mutant and WT strains; only validated strains are shown along with the corresponding WT of each experimental batch. Error bars are the S.E.M. (*n*=7).

**Figure S2.**
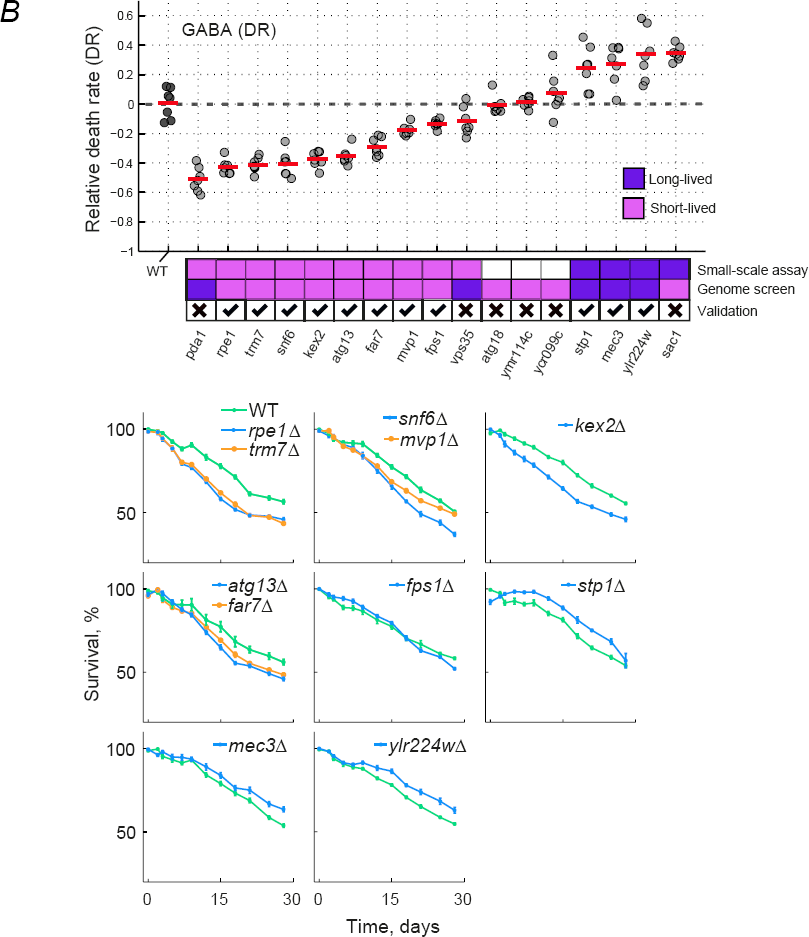
Validation of hits from the genome-wide screens. Top plot shows the relative death rates of WT and mutant strains, -1·/*n*(*r_x_/r_WT_*), aged under (**A**) non-restricted (Glutamine) and (**B**) dietary restriction (GABA) media. Death rates are from the fit of each survival curve to an exponential decay model. A CLS phenotype from the genome-wide screen was considered to be validated when the death rate of the mutant was consistent with the screen (short- or long-lived) and significantly different from that of the WT (*p*<0.05, two-tailed *T*-test). Bottom panels show the survival curves of mutant and WT strains; only validated strains are shown along with the corresponding WT of each experimental batch. Error bars are the S.E.M. (*n*=7). (Continued from previous page)

**Fig. S3.**
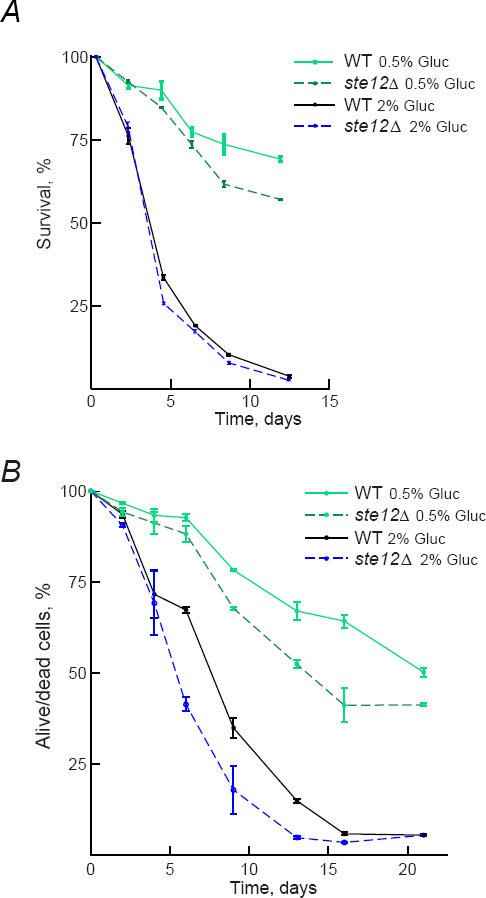
*STE12* deletion displays diminished lifespan extension measured by standard aeration conditions and alive/dead staining. (A) Survival curves obtained from single cultures aged in fully-aerated SC medium (Hu et al. 2013); WT (solid lines) or *ste12*Δ (dashed lines) strains aged under 2% or 0.5% glucose are shown. Error bars are the S.E.M. (*n*=3). (**B**) Assay of alive/dead cells as a function of time in stationary phase. Plots show the fraction of alive/dead cells in populations of WT (solid lines) or *ste12*Δ (dashed lines) strains aged under 2% or 0.5% glucose. Alive cell percentage was calculated by dividing the whole cell population in stationary phase cultures by the cells stained with Syto9 only, through time. Error bars are the S.E.M. (*n*=3).

